# Genome-Wide Association Study of Resistance to Bean Fly and Population Structure of Market Classes of Common Bean

**DOI:** 10.1101/633545

**Authors:** Pascal P. Okwiri Ojwang, Tilly Eldridge, Pilar Corredor-Moreno, Vincent Njung’e

## Abstract

Common bean (*Phaesolus vulgaris* L.) distribution across eastern, central and southern Africa region is widely driven by choice of grain types, which is affecting the genetic composition and adaptation to target production environments for biotic and abiotic constraints. Two bean fly species, *Ophiomyia spencerella* and *Ophiomyia phaseoli* are harmful insect pests of beans causing significant yield losses. Our objectives were to assess the population structure of common bean germplasm of different market classes and to identify polymorphic loci associated with resistance to *O. spencerella*. The study was carried out on a diversity panel of 284 genotypes using 9040 SNP markers. The genotypes were differentiated in to 14 distinct clusters. The mean *F*_ST_ of 0.4849, revealed major differentiation among the populations. Andean gene pool was more diverse compared to Mesoamerica gene pool which could be attributed to preference for large seeded cultivars. Multi-dimensional scaling and structure analyses revealed admixture among seed types. From genome wide association studies (GWAS), major genomic regions associated with *O. spencerella* resistance were identified on chromosome 1 (Pv01). The most significant SNP on Pv01 was aligned to gene PHAVU_001G075500g that is related to the Interleukin-1 receptor-associated kinase (IRAK) pathway, critical in regulating inherent immune responses to disease infection and insect herbivore attack. The diversity uncovered on the basis of market classes of beans and the presence of QTL regions associated with resistance to bean fly could serve as a valuable genetic resource for improvement of beans of different seed types in eastern and southern Africa region.

**Core Ideas:**

- GWAS revealed major loci associated with bean fly resistance on chromosome 01.
- New sources of resistance to bean fly were uncovered from different market classes.
- Genetic diversity uncovered is based on recent farmer preferences selection events.

## Introduction

Common bean (*Phaseolus vulgaris* L.) is the most essential grain legume for human diets, particularly in eastern and southern Africa as well as Latin America where it is a major staple food (Broughton et al., 2003; Petry et al., 2015). It is principally important as a major source of protein and micronutrients for resource disadvantaged small-holder farmers in eastern, southern and parts of central Africa (Murray-Kolb et al., 2017; Petry et al., 2015). Because of this, bean breeding programs in sub-Saharan Africa have directed efforts to develop and release new varieties for enhanced production and nutritional value that are adapted to target agro-ecologies and resilient to climate change (Blair et al., 2010a; Buruchara et al., 2011; Kamfwa et al., 2015; Kimani et al., 2001). In particular, breeders prioritize consumer preference for specific market niches of beans, which is a demand driven approach aimed at meeting the farmers’ needs for increased adoption rate (Buruchara at al., 2011).

Kenya is one of the major producers of common bean in sub-Saharan Africa and despite past breeding efforts, the on-farm production still stands at 620 kg ha^-1^ compared to the worldwide average of about 910 kg ha^−1^ (FAO 2016). The per capita consumption rate is among highest in the world ranging between 40 to 60 kg per person per year (Beebe et al., 2013; Blair et al., 2010b). Bean production is predominantly carried out under marginal conditions of drought and low soil fertility and is hampered by a multitude of biotic stresses. Therefore, developing robust breeding materials for Kenya and more widely eastern and southern Africa to address these constraints could potentially have a considerable impact on health and income of small-holder farmers (primarily women) who grow common bean (Buruchara et al., 2011).

Bean fly (Diptera: Agromyzidae), alternatively referred to as bean stem maggot (BSM) is a significant insect pest of common bean which can cause high yield losses ranging between 30 to 100 %. The most predominant species are *Ophiomyia spencerella* (Greathead) and *Ophiomyia phaseoli* (Tyron) which are adapted to high and low- to mid-altitude agro-ecologies, respectively (Songa and Ampofo, 1999). The existing management practices to control bean fly include: cultural practices, genetic diversity in the form of landraces, planting of varietal mixtures on-farm (Ssekandi et al., 2015), use of chemical and botanical pesticides and biological control. These management practices have had differing levels of effectiveness but most of them have not been scaled out to a level that can conceivably offer fortitude to the bean producers due to limited supportive scientific evidence and/or affordability for small-holder farmers. Identifying sources of genetic resistance and preserving genetic diversity will be important sustainable components in combating bean fly and increasing the resilience of the bean productions systems.

Common bean was first introduced to eastern Africa in the 17^th^ Century and cultivation started in the 19^th^ Century (Gentry 1969; Greenway, 1945). Due to this long-term cultivation and selection by farmers, eastern African region acts as a secondary centre of diversity. Long-term selection of preferred genotypes by both farmer and professional breeders affects the overall genetic conformation of modern crops (Mamidi et al., 2013; Wilkus et al., 2018). The importance of domestication process in common bean has been well documented as significant in defining the structure of genetic diversity present (Gepts and Bliss, 1988; Kwak and Gepts, 2009). However, the breeding process has led to the replacement of diverse landraces with modern varieties that have narrow genetic bases. In contrast, small-holder farmers still grow the landraces owing to their superior adaptability even though they are low yielding (Ojwang et al., 2009). These landraces are useful repositories for novel genes.

Within common bean landraces, resistance to bean fly has been reported (Kornegay and Cardona, 1991) and the genetic architecture of this resistance has been shown to be predominantly additive in nature (Ojwang et al., 2011). Few genetic studies exist on resistance to bean fly and even fewer that have exploited the genetic resources available in national gene banks. The reported studies so far concentrated on *O. phaseoli* (Abate, 1995; Ojwang et al., 2010) with little emphasis on *O. spencerella*, despite being a serious pest in higher altitude areas.

Phenotyping is an important component of selection procedure in germplasm enhancement. Breeding programs in eastern Africa currently rely on natural field infestation to phenotype breeding populations for bean fly resistance. Under field conditions, it becomes problematic to achieve uniform distribution of pest populations (Hillocks et al., 2006). Seasonal variations and environmental conditions could result in false positives when screening for resistance to bean fly. Under such circumstances, genomic selection and marker assisted breeding can be quite useful in injecting efficiency. Likewise, low cost of SNP markers has encouraged the use of genome-wide SNPs in association studies to identify useful quantitative trait loci (QTLs) for enhancement of yield, quality, biotic and abiotic stress tolerance (Mogga et al., 2018; Raman et al., 2010; Zuiderveen et al., 2016). Largely, the application of molecular markers in breeding major crops is now being embraced in sub-Saharan Africa (Ceballos et al., 2015; Edriss et al., 2017; Mogga et al., 2018). An efficient and reliable phenotyping that incorporate application of molecular markers would tap in to the genetic diversity present hence result in rapid advance in genetic improvement of common bean for biotic stress tolerance.

Previous studies on bean diversity and population structure have stratified the population in terms of geographical origin, gene pools, races as well as morphological traits (Asfaw et al., 2009; Cichy et al., 2015; Kwak and Gepts, 2009). Blair et al. 2010 reported that certain seed coat colors and patterns were associated with either Andean or Mesoamerican genepools in Rwandan accessions. Other studies using microsatellites compared accessions from different geographical regions within and across countries in eastern Africa and reported existence of minimal gene flow (Asfaw et al., 2009; Fisseha et al., 2016). The lack of correlation between genetic distance and geographical origin in common bean cultivated in Africa has been attributed to gene-flow arising from the seed systems (Kwak and Gept, 2009), which are largely non-formal and characterized by free movement of seed across regions (Wilkus et al., 2018). Generally, a limited genetic diversity exists within common bean and it is restricted to the genepools and races which is due to low levels of natural crossing arising from incompatibility among genepools (Kwak and Gepts, 2009).

Information on diversity studies focusing on market classes or seed types at molecular level for the germplasm from eastern Africa is limited. Such information would be useful for bean breeding programs to improved adoption rate by farmers (Ceccarelli and Grando, 2007; Graham and Ranalli, 1997). This is because, farmers tend to grow specific market classes (grain types) of beans (van Rheenen, 1979). In Kenya, the medium to large seeded (Andean) and in particular the red mottled (calima) and red kidneys are more popular with exception of the small seeded (Mesoamerican) red haricot (van Rheenen, 1979). The preference for large seeded has been reported in other eastern, central and southern African countries (Wilkus et al., 2018; Blair et al., 2010b). Therefore, adaptability should be combined with traits valued by small-holder farmers such as seed size, seed color, culinary qualities and cooking qualities (Hillocks et al., 2006).

The Kenya Agricultural and Livestock Research Organization (KALRO), Genetic Resources Research Institute (GeRRI) gene bank of Kenya holds over 5000 bean accessions collected from major bean growing areas across the country. The diversity of this germplasm is yet to be uncovered. The objectives of this study were to identify genomic locations associated with bean fly resistance, and to determine the genetic diversity and population structure present within the common bean germplasm from Kenya of different market classes.

## Materials and Methods

### Site Description and Plant material

The common bean genotypes were planted in the field at Egerton University, Njoro, (latitude1^°^ 15’ South and longitude 35^°^ 23’ and 36^°^ 41’ East), agronomy field 7 station of the department of Crops, Horticulture and Soils (CHS). The experimental site is situated in the highlands of the Rift Valley region, Nakuru County in Kenya. The trial was conducted for two years during the short rains July to October 2016 and the long rains May to August 2017, respectively.

The bean genotypes used for the study were largely accessions obtain from Kenya KALRO-GeRRI gene bank. To capture genetic diversity and with the help of germplasm curators at the gene bank, a representative sample was chosen based on the major gene pools (Andean and Mesoamerican) as well as range of market classes and growth habit (bush determinate and indeterminate). These considerations are particularly important and are the bases upon which farmers choose their preferred varieties. The total germplasm assembled for the study were 287. Data filtering process further reduced the number of genotypes to 284. The genotypes included, accessions, local varieties and introductions from the International Centre for Tropical Agriculture (CIAT) regional bean program, Eastern and Central Africa Bean Research Network (ECABREN) which is under the Pan-African Bean Research Alliance (PABRA). The accessions were a selection from a core collection held at GeRRI gene bank and representative of the various market classes widely cultivated in Kenya. The local varieties were recent releases from KALRO Katumani for the semi-arid region and Egerton University for the highland bean growing environments. The CIAT material had previously been screened for resistance to *Ophiomyia. phaseoli* which is a predominant species in the mid altitude areas and their inclusion in the current study was considered critical for testing them alongside the GeRRI accessions for possible dual resistance to an alternative species *O. spencerella* which is chiefly destructive in high altitude areas.

### Experimental Design and Phenotypic Data Collection

To assess the genetic variation among the genotypes for resistance to bean fly, we grew the populations over a period of two years at Egerton University, which is a high altitude environment for common bean in Kenya with a high prevalence of *O. spencerella*. The experimental design used was an alpha-lattice replicated two times, with 60 cm long single row plots, spaced 50 cm between the rows and 10 cm between plants within a row for phenotyping and to obtain leaves for DNA extraction. In year two (2017) only 256 genotypes out of the 287 were tested because some genotypes were heavily damaged by the bean fly in year one (2016) and hence had insufficient or no seed for planting. Open field screening was adopted. In each year, planting was purposefully delayed for three weeks from the onset of rainfall in order to guarantee optimal and relatively consistent spread of bean fly populations across the field for effective screening for resistance under natural open field crop infestation (Songa and Ampofo, 1999).

Due to farmer preferences for specific seed types and prevalent cultivation practices in common bean, only a few key morphological traits were considered for data collection. We collected phenotypic data on growth habit as type 1 (determinate or bush type) and type 2 (semi-determinate), seed coat color (market class) and also seed size (hundred seed weight). Genotypes weighing < 25 g per hundred seeds were considered small seeded and therefore belonged to Mesoamerican gene pool while those weighing ≥ 25 were considered large seeded hence belonging to Andean gene pool.

The data for resistance to bean fly was collected from the second week after emergence until the seventh week when all the plants had flowered. The phenotypic data collection was based on two approaches. The first approach was scoring based on a visual scale of 1-9, where 1 = immune, 2 = highly resistant, 3-4 moderately resistant, 5-6 = moderately susceptible, 7-8 = highly susceptible and 9 = extremely susceptible (Plate 1. Bean flies strike soon after seedling emergence, they oviposit their eggs on the leaves which takes about 14-18 and 28-36 days to pupate in warmer and cooler areas, respectively. We therefore scored the plants 32 days after emergence to coincide with the bean fly infestation. The second approach was based on plant mortality count which was recorded as the number of dead plants per plot. Percent mortality was then computed as the total number of dead plants resulting from bean fly attack per plot cumulative over the entire data collection period from the second week until flowering as presented below;

**Figure 1.**
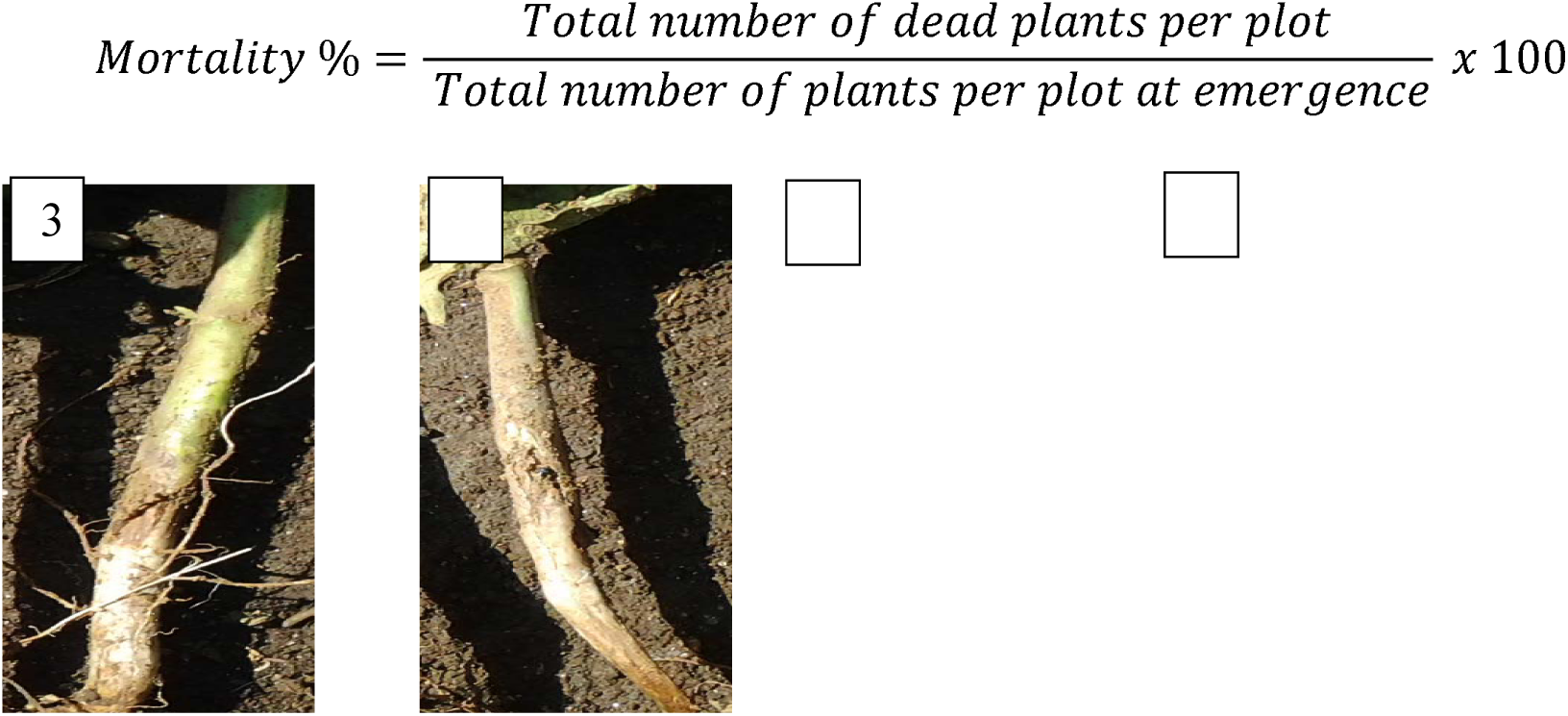
Plate 1: Damage rating based on a scale of 1-9.

### Genomic DNA Extraction and Genotyping by Sequencing (GBS)

Young leaves were collected from plants in the field and genomic DNA was extracted using the Quick-DNA^TM^ Plant/Seed-Zymo Research kit. The DNA concentrations were quantified using the Nanodrop spectrophotometer and the quality checked on 0.8% agarose gel at 5 v/cm. Approximately 30 µl of each DNA sample from all the genotypes were shipped to Diversity Array Technology (DArT) Pty Ltd. in Australia for genotyping by sequencing (GBS). A total of 15,828 SNPs were used for genotyping. Quality control (QC) was carried out and individuals with missing data > 0.4, SNPs with missing data >0.2 and minor allele frequency < 0.05 per genotype were excluded from the analysis. After filtering 9040 SNPs and 284 genotypes were retained for the analyses.

## Data Analyses

### Population Structure Analyses

The population structure among accessions was assessed using fastSTRUCTURE (Raj et al., 2014) and PLINK software (Chang et al., 2015; Purcell et al., 2007). The PLINK commands used to filter the data are given below:

1. To filter genotypes that had less than 60% genotype data, we used the PLINK command --mind 0.4.

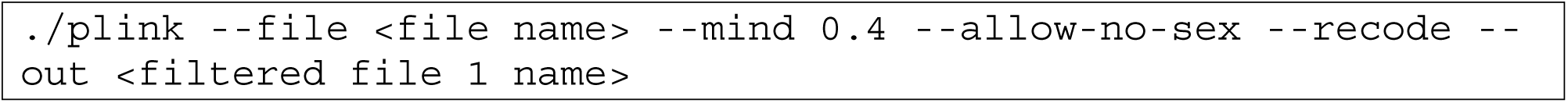
2. To filter SNPs that had < 80% genotype call rate or >20% genotype error rate, the command --geno 0.2 was used. Thus, to remove loci (SNPs) that did not have at least 80% of data from all the genotypes.

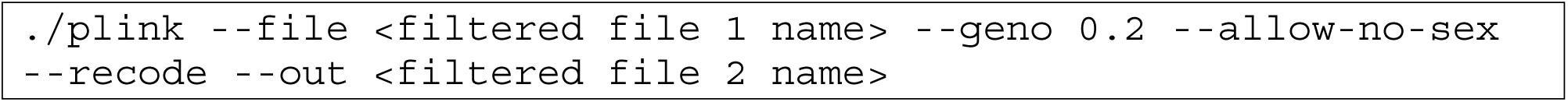
3. To filter data for rare SNPs that had > 5% minor allele frequencies, the command -- maf 0.05 was used. The command provided below removed SNPs when the second most common allele occurring in the population was present in less than 5% of the genotypes.

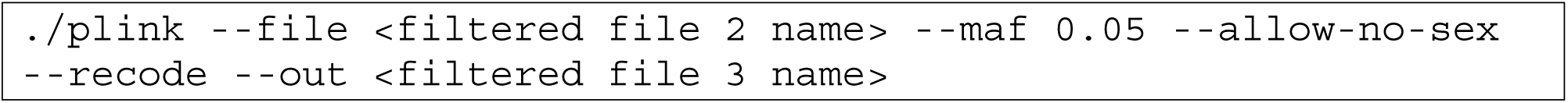

The minimum and maximum K values were set as K=2 and K=17, respectively, and this is because the population had been divided into 17 market classes (Table 1). To visualise the fastStructure results, the zipped Q files obtained from plink analysis together with their identities (ids) were uploaded to the CLUMPAK website (http://clumpak.tau.ac.il) to plot the K graph based on the admixture model.

**Table 1.** Summary of 17 market classes of bean belonging to both Mesoamerican and Andean gene pools to which the 284 genotypes were assigned.

| Market class (seed coat color) | Abbreviation§ | Number of genotypes |
| --- | --- | --- |
| **Mesoamerican gene pool (small seeded†)** |  |  |
| Buff | BF_M | 13 |
| Black | BL_M | 12 |
| Cranberry | CB_M | 9 |
| *Mwitemania* (pinto bean) | MW_M | 6 |
| Red (small red or haricot) | RD_M | 19 |
| *Rose coco* | RS_M | 2 |
| White | WT_M | 42 |
| **Andean gene pool (large seeded‡)** |  |  |
| Buff | BF_A | 29 |
| Black | BL_A | 3 |
| Cranberry | CB_A | 33 |
| *Mwezi moja* (variegated grey with cream or purple background) | MM_A | 5 |
| *Mwitemania* (pinto bean) | MW_A | 3 |
| Purple | PP_A | 6 |
| Red | RD_A | 39 |
| Red mottled | RM_A | 20 |
| *Rose coco* | RS_A | 32 |
| White | WT_A | 11 |
| **Total** |  | **284** |

† Small seeded $\geq 25g$ 100 seed weight<sup>-1</sup>.
‡ Large seeded $> 25g$ 100 seed weight<sup>-1</sup>.
§ The Mesoamerican gene pool was represented by 103 genotypes and the Andean by 181 genotypes.

The molecular data and both individual and market class (population) data were used to perform multi-dimensional scaling (MDS) in plink. The squared distances were determined and R package was used for visualization (R Core Team, 2018 Library Ape). The mean fixation statistics (F_ST_) (Weir and Cockerham, 1984) was also determined.

### Phenotypic Data Analyses

Phenotypic data were subjected to residual maximum likelihood (REML) analysis in GenStat version 15 statistical package. The phenotypic data were tested for the assumptions of analysis of variance (normality, homogeneity and additivity). Data under study were percentages and therefore were arcsine-square root (angular) transformed before analysis. The statistical model used to obtain the best linear unbiased predictors (BLUPs) is provided below:

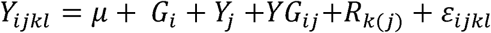

Where *Y_ijkl_* is the phenotype (percent plant mortality) of the *i^th^* genotype in the *j^th^* year and the *k^th^* replicate, *G_i_* is the random effect of the *i^th^* genotype, *Y_j_* the random effect of the *j^th^* year, *YG_ij_* is the random effect of the year x genotype interaction, *R_k_* is the fixed effect of the *k^th^* replicate within the *j^th^*year, and *ε_ijkl_* is the random error term.

### Association Studies

R-based Genome Association and Prediction Integrated Tool (GAPIT) software package was used for genome-wide association studies (GWAS) analysis (Lipka et al., 2012). In order to eliminate two sources of false positives mainly the population structure and familial relatedness, a unified linear mixed model (MLM) was used as given below:

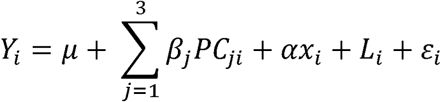

Where *Y_i_* is the phenotype of the *i^th^* individual, *µ* is the grand mean, *B_j_ PC_ji_* is the fixed effect accounting for the population structure considering the first three principal components (PCs), *α* is the marker effect, *x_i_* is the observed SNP alleles of the *i^th^* individual, *L_i_* random effects accounting for familial relatedness and *ε_i_* is the random error term. The threshold used for significant SNP markers for percent mortality was the Bonferonni corrected *P* = 1.106 x 10^−5^ (for α = 0.05 and 9040 SNPs).

## Results

### Population Differentiation and Structure

We assembled 287 common bean (*Phaseolus vulgaris* L.) germplasm belonging to different market classes to determine the population structure. Among these genotypes, 269 were accessions from the Kenya Agricultural and Livestock Research Organization, Genetic Resources Research Institute gene bank (KALRO-GeRRI) primarily to capture selection preferences of Kenyan and largely eastern and southern African farmers. The 269 genotypes were selected on the basis of seed coat color (market class), seed size or gene pool (small seeded mainly Mesoamerican and medium to large seed mainly the Andean gene pool) and growth habit (determinate and indeterminate growth). The accessions from KALRO-GeRRI, classed as landraces, also captured local adaptation to marginal production environments. Additionally, 10 genotypes (accessions and improved varieties) from Eastern and Central Africa bean Research Network (ECABREN) which had been reported to be resistant to *O. phaseoli* bean fly species, were included. The control genotypes were 5 elite local varieties recently released from KALRO Katumani research station for the semi-arid regions and 3 from Egerton University for the highland regions and all of them belonging to preferred market classes. The population differentiation was therefore founded on the market class (seed coat color), gene pools (Table 1) which is the basis for farmer and breeder preferences of bean varieties. It was difficult to use geographical distribution since over 80% of the accessions lacked passport data. Together, this panel of genotypes allowed us to capture common bean gene pools, major market classes of beans and tolerance to bean fly.

To assess the structure of the population, structure analysis, multidimensional scaling (MDS) were carried out and fixation statistics (F_ST_) estimated using PLINK, fastSTRUCTURE and R. Details of the statistical approaches adopted can be found in the materials and methods section. The results depicted by fastSTRUCTURE analysis ascertained the presence of two major bean gene pools of common bean among the market class populations (Figure 1). At K = 2, the 17 populations were classified into two genetic groups: Andean and Mesoamerican gene pools, with the buff and cranberry Andean genotypes showing slightly more but generally low admixture compared to the rest of the Andean genotypes. Among the Mesoamerican genotypes it was the white seeded genotypes that showed relatively more admixture than the other genotypes even though it was comparatively low. At K = 3, the Andean genotypes showed more differentiation again with the buff and cranberry being separated in to two sub-groups with low admixture which generally remained the same up to K=6 except for minimal separation at K = 4 and K=6, however, no meaningful conclusion could be drawn from this differentiation. On the other hand, the Mesoamerican genotypes showed no differentiation from K = 2 to K = 4, but a visible separation was observed at K = 5 of which a sub-grouping comprising of buff, cranberry and *rose coco* market classes was created with the white forming its own separate cluster. The pattern for the Mesoamerican gene pool remained relatively the same from K = 5 up to K = 14. However, at K=7 the Andean gene pool had a distinct differentiation where buff, black, cranberry purple and red mottled formed a sub-group from the rest of the market classes. From K = 9 to K = 14, the Andean gene pool maintained the sub-grouping except for K = 10 which appeared to lump all the market classes together. Overall, the model complexity that maximizes marginal likelihood was K = 1 while the model components used to explain structure were K=14 by the fastStructure procedure in PLINK software. Similarly, a high mean *F*_ST_ of 0.4849 was estimated as the level of differentiation among the populations. From this population structure analysis, the Andean gene pool appeared more diverse compared to Mesoamerican gene pool.

**Figure 1.**
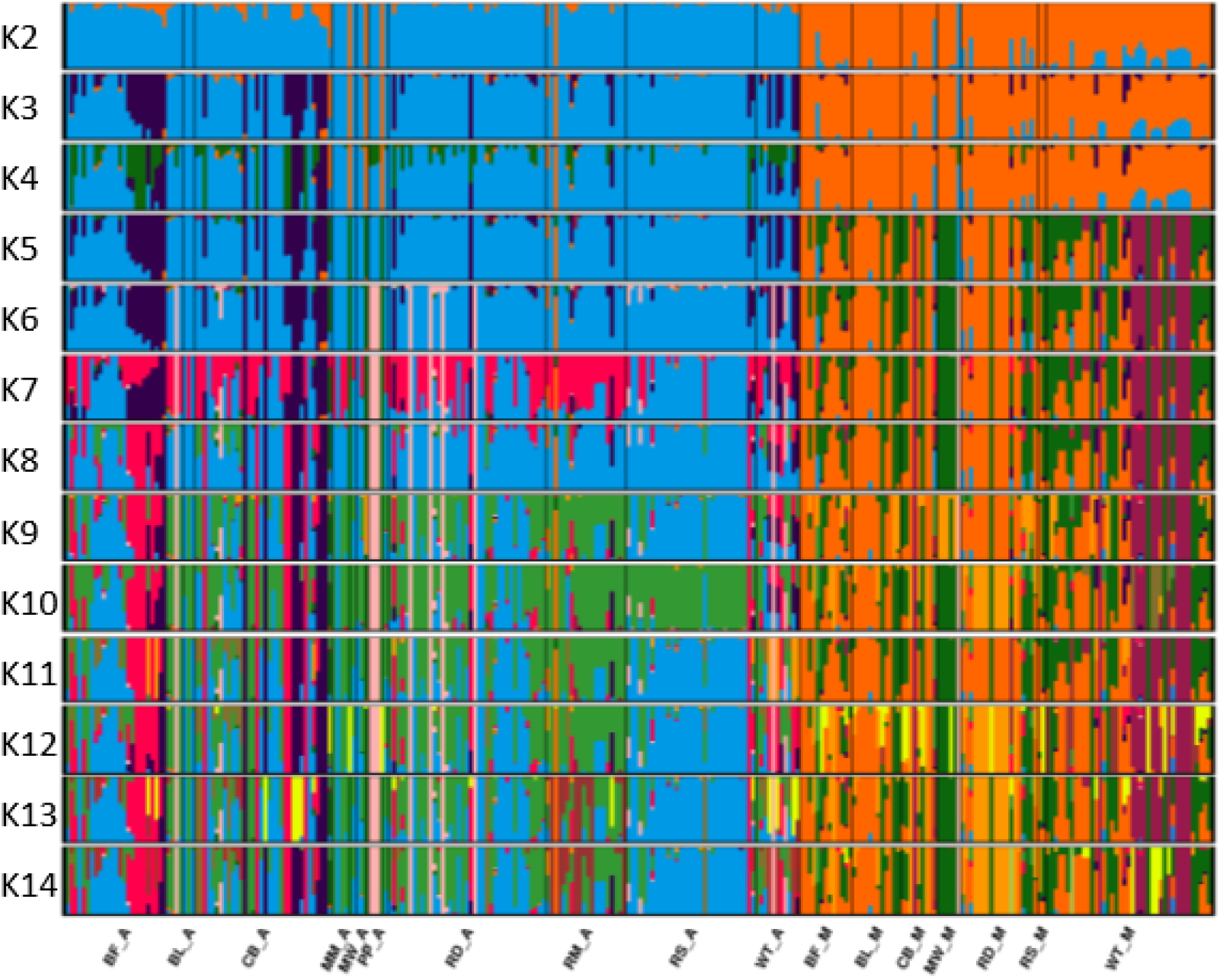
Genetic kinship of 284 common bean genotypes based on 9040 genome-wide SNP markers and analyzed by plink and Structure plot bar graph generated in CLUMPAK program for K = 2 - 14. Market class populations were classed by seed coat color (BF, buff; BL, black; CB, cranberry; MM, *mwezi moja* or variegated; MW, *mwitemania* or Pinto type; PP, purple; RD, red; RM, red mottled or calima type; RS, *rose coco*; WT, white) and the gene pools, A and M Andean and Mesoamerican, respectively.

We further assessed the structure of common bean market class population using a pairwise *F*_ST_ matrix estimates. Following the analysis, evidence of significant differentiation between the Andean and Mesoamerican gene pools was obtained (Figure 2). The Andean gene pool being presented by 10 market classes were differentiated in to 3 main clusters. The first major cluster had four market classes which were the navy beans (white seeded) (WT_A), the red mottled seed type (RM_A), cranberry (CB_A) and buff (BF_A). The second cluster was differentiated in to *mwitemania* (MW_A; a pinto type) and black seed coat (BL_A) market classes. The rose coco (RS_A) and the large red seeded type (RD_A) formed the third cluster, while purple (PP_A) and *mwezi moja* (MM_A; a variegated seed color with grey as main color with pink or cream background) formed the fourth cluster. Conversely, the accessions that belonged to the Mesoamerican gene pool were all lumped in to one major cluster having all the seven market classes viz; cranberry, buff, red, black, *rose coco*, white and *mwitemania* seed types. However, cranberry and buff seed types appeared closely related. Generally, the genetic structure of bean market classes belonging to the Andean gene pool showed a broader diversity compared to the Mesoamerican gene pool.

**Figure 2.**
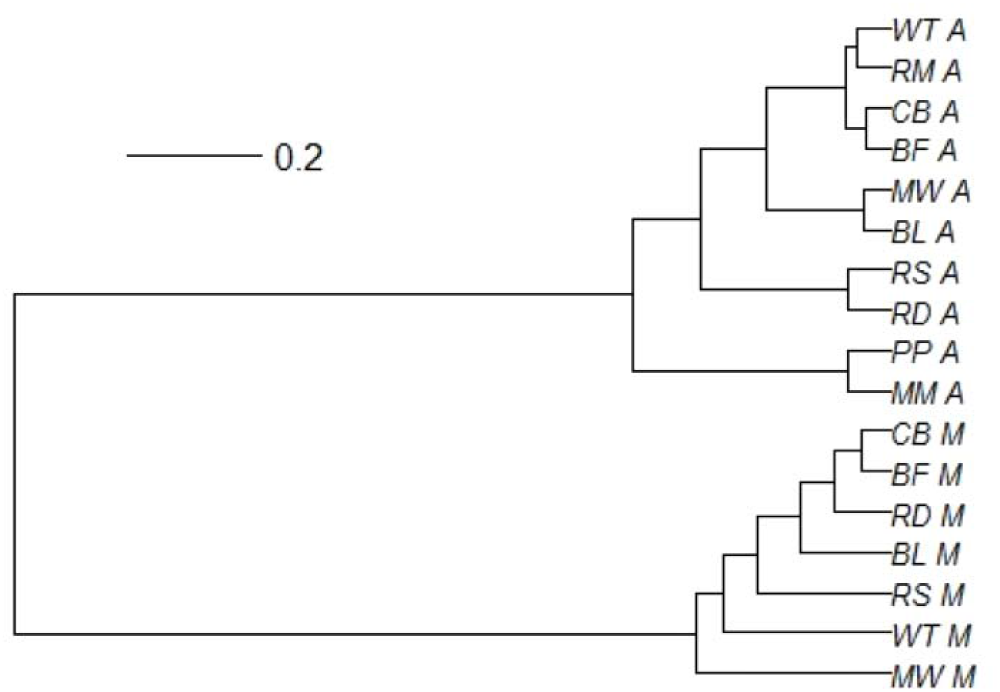
Pairwise *F*_ST_ cluster dendrogram for 17 common bean market classes belonging to Andean and Mesoamerican gene pools. The letters A and M stand for Andean and Mesoamerican gene pools, respectively while the abbreviations are for the market class populations classified by seed coat color (BF, buff; BL, black; CB, cranberry; MM, *mwezi moja* or variegated; MW, *mwitemania* or Pinto type; PP, purple; RD, red; RM, red mottled or calima type; RS, *rose coco*; WT, white).

To take a further look at population structure we used the multi-dimensional scaling (MDS) plot to show the stratification of the accessions (Figure 3). For the first two coordinates (C1 and C2), the accessions were generally separated in to Andean and Mesoamerican gene pools. The pattern of differentiation was in agreement with the Structure analysis and *F_ST_* cluster dendrogram which clearly showed that the large seeded (Andean) seed types; rose coco, black, buff and cranberry tended to cluster together. Contrary, the small seeded (Mesoamerican) did not show any clear pattern of clustering. In all, our population structure analysis results revealed more differentiation among the Andean gene pool compared to their Mesoamerican counterparts.

**Figure 3.**
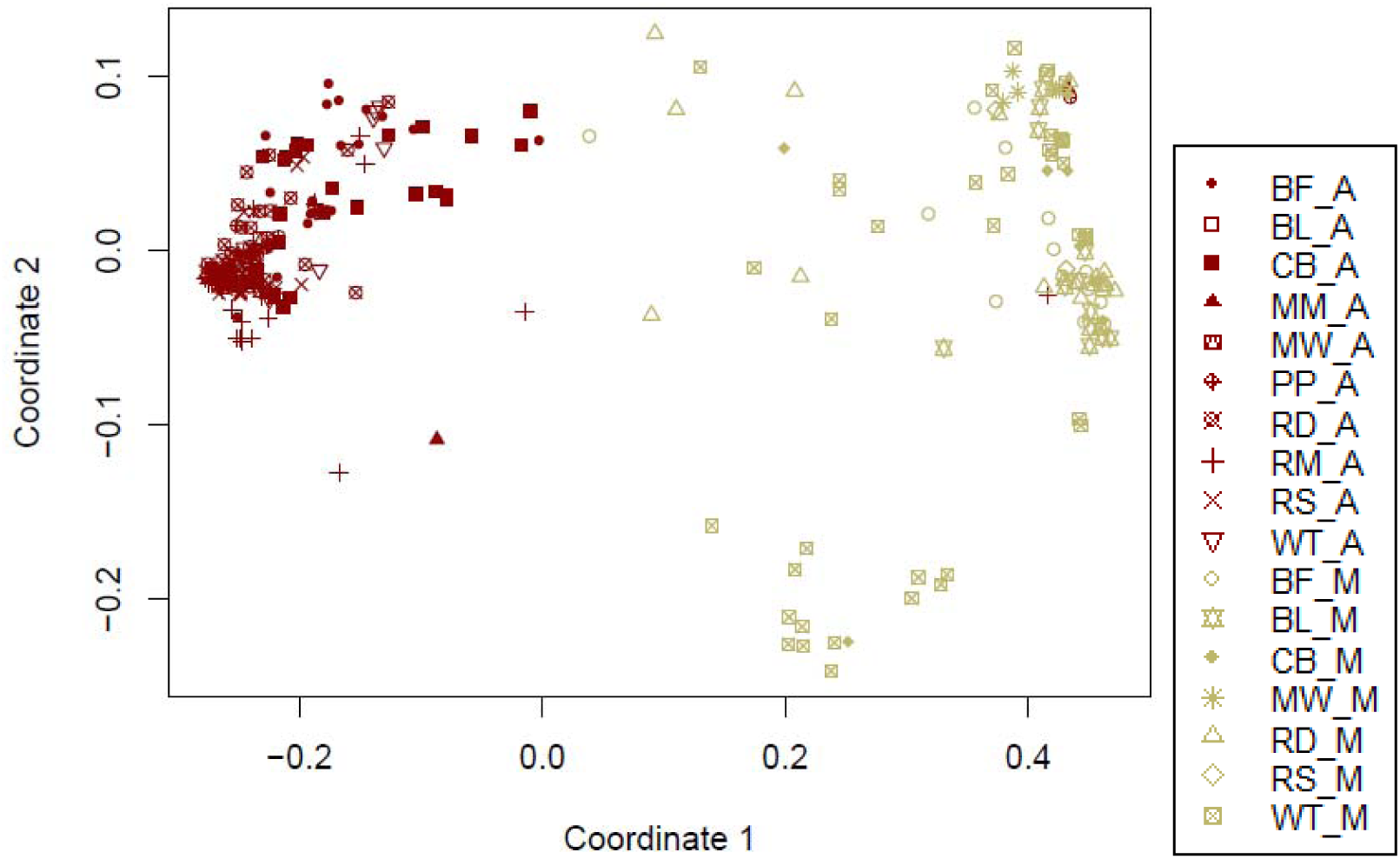
Multi-dimensional scaling (MDS) for 17 bean market class populations. The letter suffixes A and M placed on the figure legend after the underscores stand for Andean and Mesoamerican gene pools while the abbreviations are for the market class populations representing seed coat color (BF, buff; BL, black; CB, cranberry; MM, *mwezi moja* or variegated; MW, *mwitemania* or Pinto type; PP, purple; RD, red; RM, red mottled or calima type; RS, *rose coco*; WT, white).

### Phenotypic evaluation for bean fly resistance

We grew the common bean diversity panel in highland conditions and phenotyped them for bean fly resistance over two cropping seasons. The results showed phenotypic variation among common bean genotypes based on visual scoring (Figure 4) and percent plant mortality (Figure 5). Plant stem damage was score only once, 32 days after emergence while plant mortality was cumulative from 14 days after emergence up to anthesis (a period of 6 weeks). Genotypes that had a score of <3 were considered resistant and those that had >7 were susceptible. On the other hand, genotypes that showed < 30% mortality by anthesis were considered resistant and those that showed > 70% mortality by anthesis were considered susceptible. Comparing data obtained from these two approaches, percentage mortality was a more reliable approach. This is owing to the ability of some bean genotypes to recover by developing adventitious roots above the damaged area hence turn out to be tolerant.

**Figure 4.**
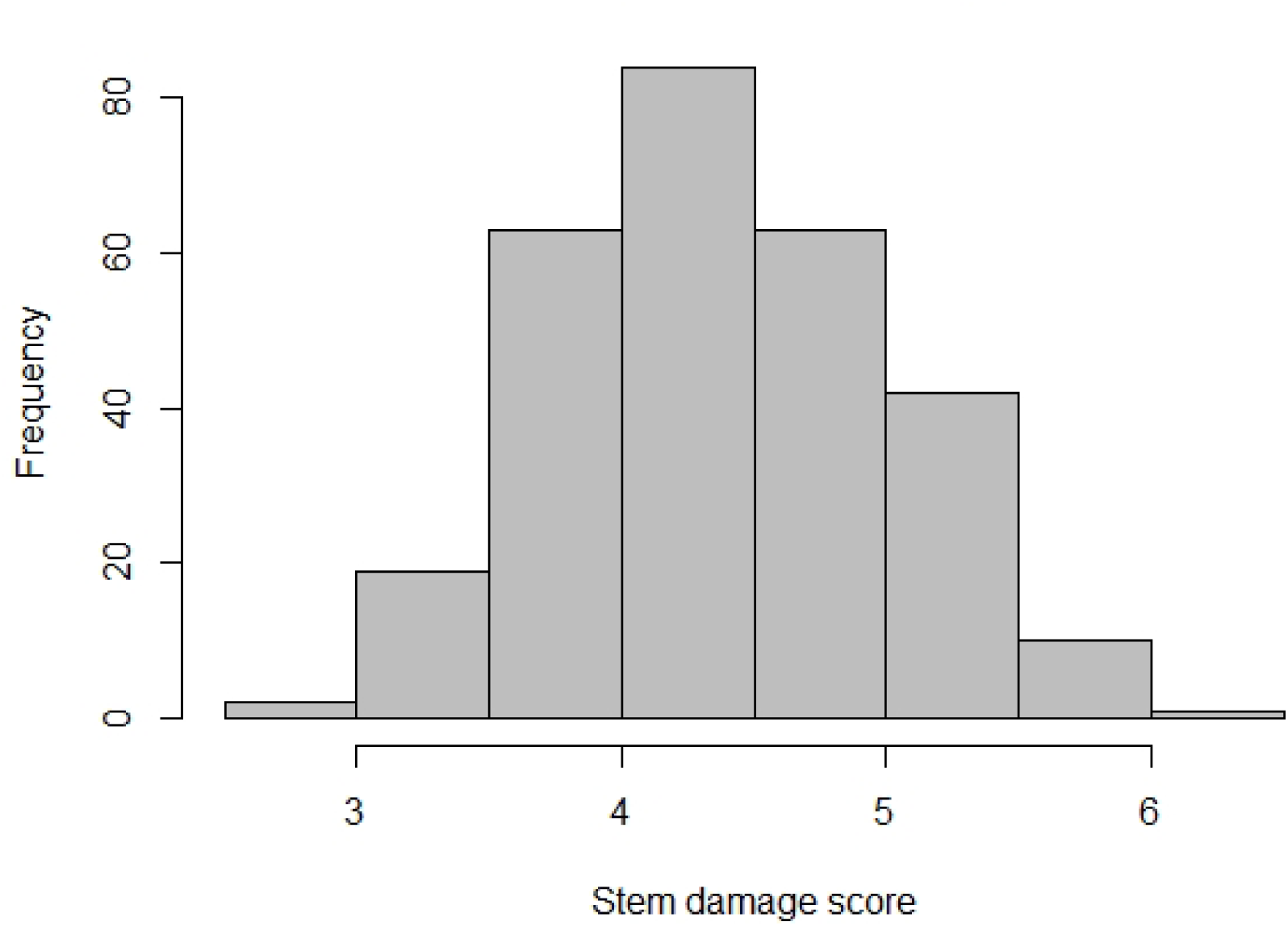
The frequency distribution of stem damage score of 284 genotypes evaluated in this study under field conditions combined over the two years, 2016 and 2017.

**Figure 5.**
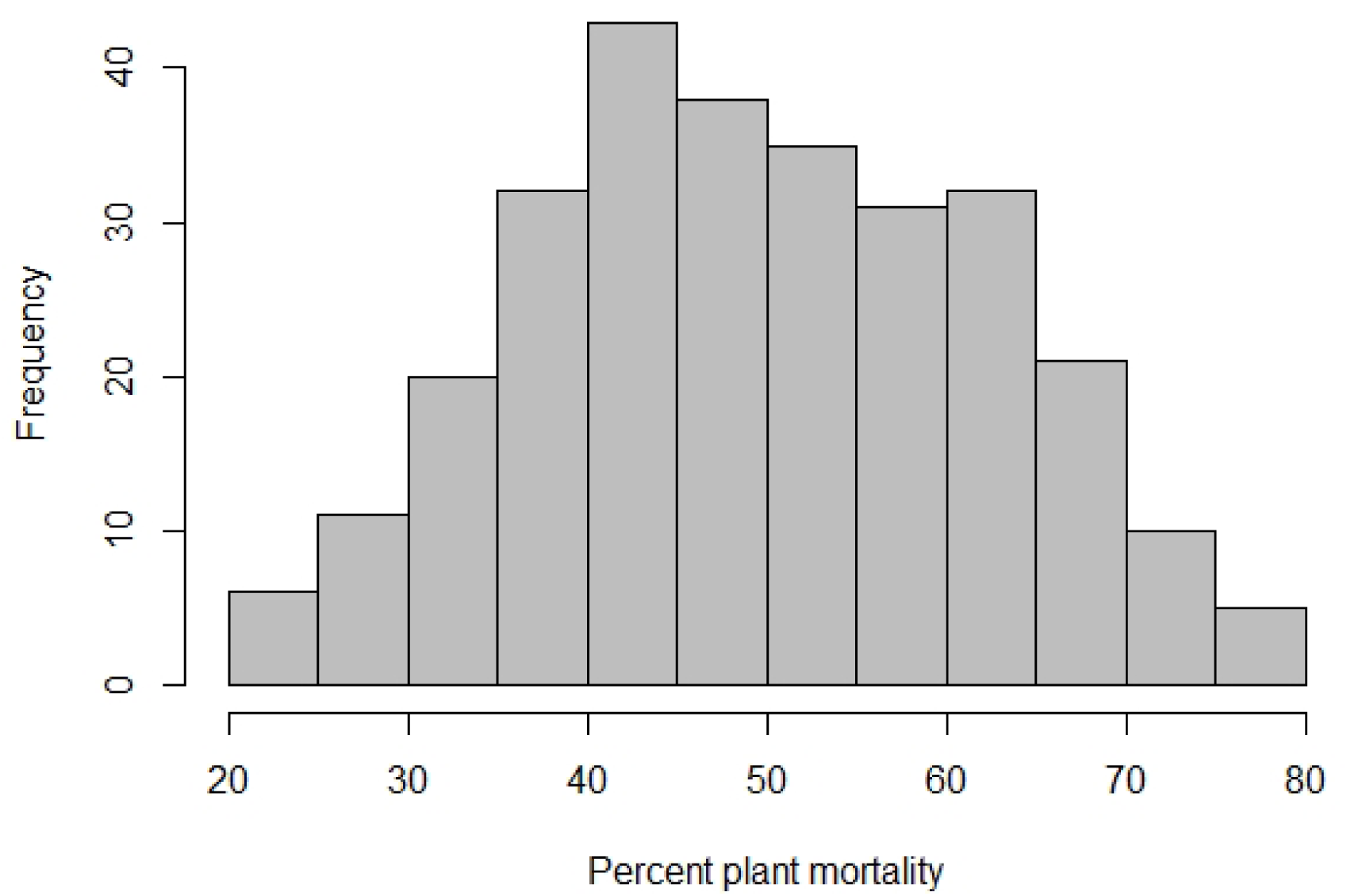
The frequency distribution of percent plant mortality of 284 genotypes evaluated in this study under field conditions combined over the two years, 2016 and 2017. Genotypes with a percent mortality of < 30% were considered resistant.

Among the germplasm, 17 accessions exhibited high resistance with a mortality rate of <30% (Table 2). Out of the resistant genotypes, only one was from CIAT (G1212) the rest were accessions from GeRRI (Table 2). We observed that the most resistant accessions proportionately belonged to both gene pools and wide-ranging market classes. Therefore, we did not observe a link between a given population and the pattern of resistance.

**Table 2.** List of accessions which showed < 30% mortality (resistance) to *Ophiomyia spenceralla* species of bean fly in the diversity panel over the two years of field evaluation at Egerton University agronomy field 7 station in 2016 and 2017.

| Landraces | Source | Gene pool | Market class (seed type) | Growth habit† | Percent plant mortality‡ |
| --- | --- | --- | --- | --- | --- |
| GBK-032833 | GeRRI | Andean | Red mottled | Type 1 | 20.62 |
| GBK-035141 | GeRRI | Mesoamerican | Black | Type 2 | 21.09 |
| GBK-035104 | GeRRI | Andean | Red | Type 2 | 22.91 |
| GBK-032803 | GeRRI | Mesoamerican | Red | Type 2 | 22.91 |
| GBK-028030 | GeRRI | Andean | Variegated (*Mwezi moja*) | Type 1 | 23.32 |
| GBK-035133 | GeRRI | Mesoamerican | Buff | Type 1 | 23.82 |
| GBK-035106 | GeRRI | Mesoamerican | Red | Type 2 | 26.73 |
| GBK-032625 | GeRRI | Mesoamerican | Red | Type 2 | 27.14 |
| GBK-032865 | GeRRI | Mesoamerican | Buff | Type 1 | 27.99 |
| GBK-032897 | GeRRI | Andean | Purple | Type 2 | 27.99 |
| GBK-035233 | GeRRI | Mesoamerican | White | Type 1 | 28.37 |
| GBK-035046 | GeRRI | Andean | *Rose coco* | Type 1 | 29.01 |
| GBK-027810 | GeRRI | Andean | Cranberry | Type 1 | 29.12 |
| GBK-035069 | GeRRI | Andean | *Rose coco* | Type 1 | 29.14 |
| GBK-035651 | GeRRI | Andean | Cranberry | Type 2 | 29.32 |
| G1212 | CIAT | Mesoamerican | Black | Type 2 | 29.58 |
| GBK-035103 | GeRRI | Mesoamerican | Red | Type 2 | 29.64 |

† Type = Bush (indeterminate growth); Type 2 = Semi-determinate.
‡ Back transformed means ( $\sin(x)^2 \times 100$ ).

### Association Studies for Resistance to Bean Fly (*Ophiomyia spencerella)*

Genotyping by sequencing using Diversity Arrays Technologies sequencing platform (DarT-seq) was done for all the genotypes and after filtering using PLINK software (Chang et al., 2015) we retained 9040 SNPs. The filtered tped and tfam file formats from PLINK output was converted to hapmap file format for association analysis. We then used Genome Association and Prediction Integrated Tool (GAPIT) in R package (Lipka, 2012) for GWAS analysis. From the results, we depicted the existence of a region in chromososome 1 (Pv01) where resistance to bean fly appear to be exclusively positioned within common bean populations (Figure 6). Overall, five SNPs were significantly associated with resistance to *Ophiomyia spencerella*. The most significant SNP (SNP372_8196616) had a *P*-value = 3.64 x 10^−7^ which explained 18.64 % of the variability with a false discovery rate (FDR) *P*-value = 0.001656 (Table 3) and fell in the range of 11-12 x 10^−6^ base pairs (Figure 6). The second most significant SNP (SNP373_3376280) had a *P*-value = 3.68 x 10^−7^ with 18.63 % variation explained followed by SNP379_3377589 (*P*-value = 6.34 x 10^−7^) with 18.2 % variation explained. The remaining significant SNPs explained 17.19 and 16.98% variation, respectively. Thus, we found 5 highly significant SNPs in the same linkage group that were effective in conditioning resistance to bean fly.

**Figure 6.**
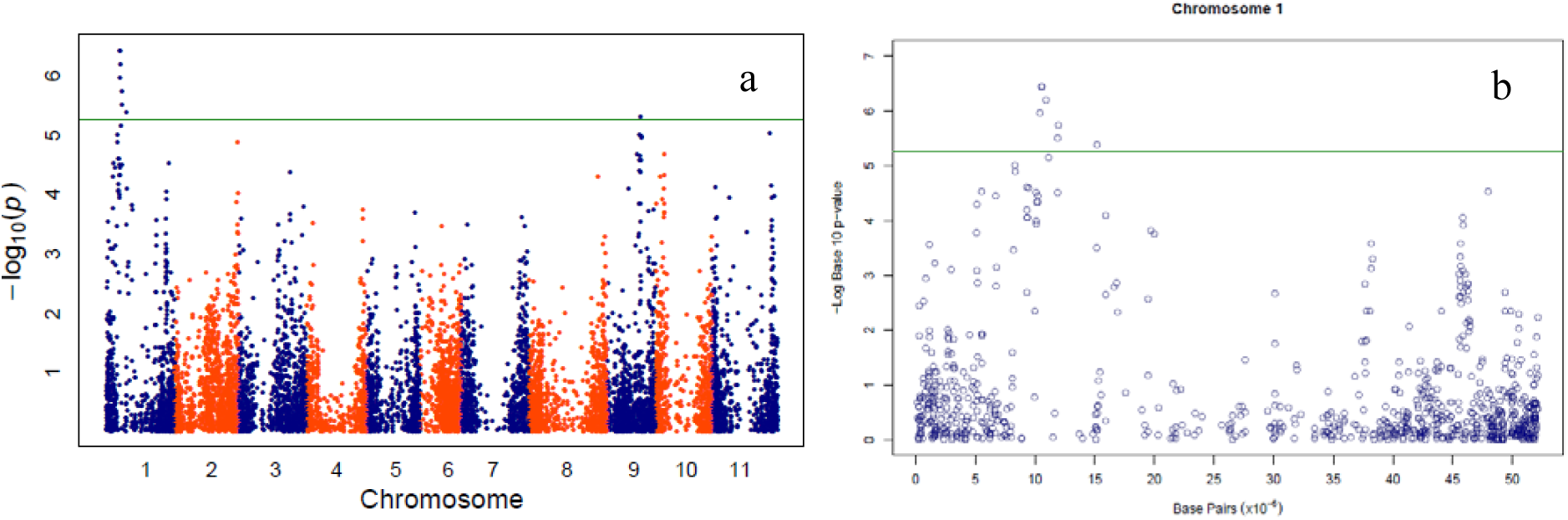
Genome-wise (a) and chromosome 1 (b) Manhattan plots showing significant SNPs and *P*-values from GWAS analysis. Green line on Manhattan plots are the threshold significance based on the Bonferonni correction at *P* = 1.106 x 10^−5^ (for α = 0.05 and 9040 SNPs).

**Table 3.**
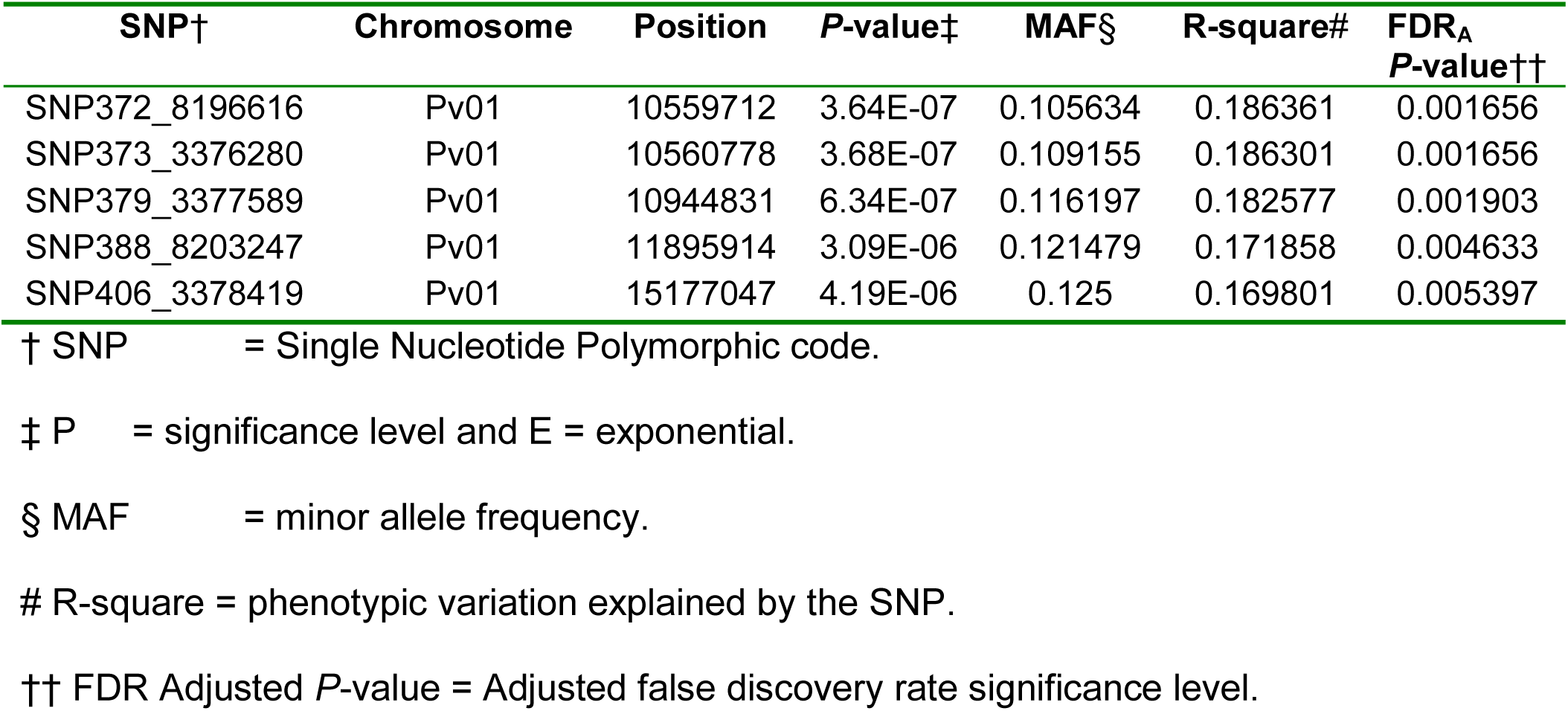
List of the most significant SNPs associated with resistance to *Ophiomyia spenceralla* species of bean fly measured on 284 common bean genotypes.

| SNP† | Chromosome | Position | P-value‡ | MAF§ | R-square# | FDR<sub>A</sub><br>P-value†† |
| --- | --- | --- | --- | --- | --- | --- |
| SNP372_8196616 | Pv01 | 10559712 | 3.64E-07 | 0.105634 | 0.186361 | 0.001656 |
| SNP373_3376280 | Pv01 | 10560778 | 3.68E-07 | 0.109155 | 0.186301 | 0.001656 |
| SNP379_3377589 | Pv01 | 10944831 | 6.34E-07 | 0.116197 | 0.182577 | 0.001903 |
| SNP388_8203247 | Pv01 | 11895914 | 3.09E-06 | 0.121479 | 0.171858 | 0.004633 |
| SNP406_3378419 | Pv01 | 15177047 | 4.19E-06 | 0.125 | 0.169801 | 0.005397 |

† SNP = Single Nucleotide Polymorphic code.
‡ P = significance level and E = exponential.
§ MAF = minor allele frequency.

### Discussion

Using genome-wide SNPs we uncovered the presence of considerable level of genetic diversity among the accessions (from Kenya). Consistent with previous reports, the accessions used in this study were grouped in to Andean and Mesoamerican geneools (Cichy et al., 2015; Gepts and Bliss, 1988; Kwak et al., 2009; Shi et al., 2011). Based on the diversity panel of market classes of beans commonly grown in Kenya, our results corroborated previous reports of high molecular and phenotypic diversity among common bean germplasm from different geographic regions within or among countries in eastern Africa (Asfaw et al., 2009; Fisseha et al., 2016).

Phenotypic variability observed in this study enabled us to place the germplasm in to 17 market classes primarily on basis of seed coat color and seed size. However, there was an indication of admixture among some market class populations from the fixation statistics where the model components used to explain structure in data equaled 14. McClean (2002) reported variation in seed coat color, pattern and shape in common bean. Although farmers in Kenya largely grow large seeded market classes of beans, small seeded types may also be preferred depending on specific attributes (van Rheenen, 1979). In line with these farmers’ preferences, the breeding programs in the eastern Africa region through participatory breeding and variety selection have also transformed and are currently focusing on combining genes for grain type with major biotic and abiotic constraints as multiple trait introgression approach (Buruchara et al., 2011). To contribute to this goal, this study has used farmers’ priorities for selected market classes as a basis for understanding the population structure and diversity of common bean populations in Kenya.

A high *F*_ST_ value of 0.4849 obtained from our study suggests a recent origin or a selection event (Kwak and Gepts, 2009). The selection is most likely associated with preference for market classes as well as growth habit of which the bush and semi-determinate types are predominant due to their compatibility to the intercropping systems, a common practice by small-holder farmer in the eastern Africa region. Farmer selection and preference for propagating a selected market classes of beans appear to contribute to the pattern of genetic diversity. This is influenced by consumption and commercialization of beans. Apart from farmer selection practices, low levels of natural crossing between the Andean and Mesoamerican gene pools is probably another source of the differentiation. The informal seed network for beans in Kenya has resulted in random distribution of bean seed and therefore classification of material based on geographic distribution may fail to reflect the true differentiation. Our results suggest that the bottleneck of domestication from the primary centres at the time of introduction is important in explaining the diversity present in common bean germplasm in Kenya, which is in agreement with earlier reports (Burle et al., 2010; Mamidi et al., 2013).

Domesticated common beans tend to have lower genetic diversity and higher *F*_ST_ (Kwak and Gepts, 2009). Considering the pairwise *F*_ST_ values, the molecular analysis revealed high differentiation between gene pools with a low level of gene flow between them. Nevertheless, evidence of some level of inter-gene pool admixture was observed from the current study even though it was low. Conversely, the gene flow between the market classes within gene pools was relatively high. Limited admixture was observed between Andean and Mesoamerican gene pools (Burle et al., 2010). Recently, Lobaton et al. 2018 reported the presence of inter-introgression which demonstrated the admixture events in the breeding history of common bean.

In spite of their importance, the Andean gene pool has been previously reported to be less diverse compared to the Mesoamerican gene pool (Burle et al., 2010; McClean et al., 2012). However, in our study, the Andean gene pool was more diverse compared to the Mesoamerican gene pool. This was expected because of the cultivation practices that tend to promote the large seeded types. Beans from this gene pool are quite important to nutrition and income generation among small-holder farmers in Kenya. Therefore, characterization of local germplasm to assess diversity is key towards improvement of food security in Kenya.

The accessions used in the current study are potentially a valuable genetic resource for improvement of the predominant market classes of common bean for diverse production environments in Kenya and potentially in eastern, central and southern Africa. For example, genotype G1212 appeared to possess resistance to both *O. phaseoli* and *O. spencerella*, having been identified from an earlier study to be resistant to the former species of bean fly. Identification of new sources of genes for inclusion in the bean improvement programs as a strategy for multiple trait integration to combine market class preference, adaptive traits (biotic and abiotic stress tolerance) and yield potential is key to enhancement of bean productivity.

Resistance to insect pest has not been extensively studied compared to disease resistance and thus has led to limited availability of information for breeding programs aiming at introgression of insect resistance traits in to elite cultivars. This has resulted in few varieties with insect resistance being released (Miklas et al., 2006). Host plant resistance is a useful component of common bean field pest management with known benefits of minimizing use of chemical pesticides and increasing on-farm yield at lower cost of production. Previous studies pointed out that key mechanisms involved in bean fly resistance are antixenosis and antibiosis (Cardona and Kornegay, 1999) which are controlled by quantitative trait loci with several genes contributing to the resistance (additive gene effects) (Ojwang et al., 2011). For improved efficiency in breeding, application of molecular markers is crucial for the deployment of novel resistance genes. In the current study, 17 accessions were identified to hold bean fly resistance alleles which could be useful for future breeding efforts.

Genome-Wide Association Studies (GWAS) revealed major loci controlling bean fly (*Ophiomyia spencerella*) on common bean chromosome 1 (Pv01). The genomic region associated with the most significant SNP (SNP372_8196616) was investigated for possible putative genes and the results on BLASTx aligned 100% to a hypothetical protein gene PHAVU_001G075500g on chromosome 1 (pv01). A follow up on this gene on BLASTn matched the gene to Interleukin-1 receptor-associated kinase (IRAK). This protein kinase super-family is principally composed of the catalytic domains of serine-threonine-specific and tyrosine-specific protein kinases. The IRAKs are involved in the interleukin-1 (IL-1) signaling pathways, and are thus critical in regulating inherent immune responses (Li et al., 1997). These proteins include Leucine-rich repeat receptor-like kinases (LRR-RLKs) such as the Arabidopsis thaliana BAK1 which functions in BR (brassinosteroid)-regulated plant development and in pathways involved in plant resistance to pathogen infection and herbivore attack (Dievart and Clark, 2004). BAK1 regulates BR-dependent developmental responses, and equally controls pathways involved in resistance to pathogen infection and herbivore attack, even though the functional mechanisms involved in the two biotic stresses differ (Yang et al., 2011). Potentially, this is the first identification of a gene that is associated with bean fly resistance and provides a basis for future characterization together with a strongly associated molecular marker for introgression.

The quantitative trait loci (QTLs) identified in this study should be validated. If found to be stable and expressed in different genetic backgrounds of common bean market classes, they can be used as potential candidates for a marker-assisted breeding program for bean fly resistance. If the markers do not account for sufficient genetic variation, marker assisted selection (MAS) may not be the most efficient method (Meuwissen et al. 2001). Further, since phenotyping for bean fly resistance relies heavily on plant mortality data which is destruction of individual plants by insect, genomic selection (GS) offers a more robust option. The next step will be to develop a training population set of individuals with marker information and high-quality estimates of breeding values that can be phenotyped and genotyped to generate a training genomic selection (GS) model. The model can be used to predict trait values for a breeding test set. This could help in speeding up the generation time required to develop a new variety carrying bean fly resistance.

## Conclusions

This study has uncovered great genetic diversity present in the common bean germplasm collections conserved at the Kenya Agricultural and Livestock Research Institute, Genetic Resources Research Institute (KALRO-GeRRI), regional and local breeding programs. Further, the GWAS discovered QTL regions on Pv01 that were significantly associated with bean fly resistance. Some of these QTLs could potentially be employed in Marker Assisted Breeding to speed up the genetic gains in breeding for resistance to bean fly herbivory. However, additional investigations will be required using segregating populations at the significant SNP loci in order to validate the QTLs and to determine their practicality in application in the breeding programs.

## Abbreviations

BAK1: brassinosteroid insensitive 1-associated receptor kinase 1
BLASTn: basic local alignment search tool for nucleotide
BLASTx: basic local alignment search tool for translated nucleotide
BR: brassinosteroid
BSM: bean stem maggot
CIAT: International Centre for Tropical Agriculture
DArT: Diversity Array Technology
ECABREN: Eastern and Central Africa Bean Research Network
FAOSTAT: Food and Agricultural Organization statistics
FDR: false discovery rate
GBS: Genotyping by sequencing
GeRRI: Genetic Resources Research Institute
GS: genomic selection
GWAS: genome-wide association study
IRAK: Interleukin-1 receptor-associated kinase
KALRO: Kenya Agricultural and Livestock Research Organization
MAF: minor allele frequency
MDS: multi-dimensional scaling
MLM: mixed linear model
PABRA: Pan-African Bean Research Alliance
PC: principal component
Pv: Phaseolus vulgaris
REML: residual maximum likelihood
QTL: quantitative trait loci
SNP: single nucleotide polymorphism.

## Acknowledgments

This work was fully supported by BecA-ILRI Hub through the Africa Biosciences Challenge Fund (ABCF) program. The ABCF program is funded by the Australian Department for Foreign Affairs and Trade (DFAT) through the BecA-CSIRO partnership; the Syngenta Foundation for Sustainable Agriculture (SFSA); the Bill & Melinda Gates Foundation (BMGF); the UK Department for International Development (DFID) and the Swedish International Development Cooperation agency (Sida). The authors express their gratitude to the UK Biotechnology and Biological Sciences Research Council, Global Challenges Research Fund for funding Tilly Eldridge’s and Pillar Corredor-Moreno’s participation in the project. We thank, the Kenya Agricultural and Livestock Research Organization (KALRO), Genetic Resources Research Institute (GeRRI) and the International Centre for Tropical Agriculture; Eastern and Central Africa Bean Research Network (CIAT-ECABREN) providing the germplasm. Egerton University is acknowledged for providing research experimental fields and logistical support. David Ongige is recognized for assisting with field management.

## Author Contributions

Conceived and designed the experiments: Ojwang, P.P.O., and T. Eldridge. Performed the phenotyping: Ojwang, P.P.O. Analyzed the data: Ojwang, P.P.O and Corredor-Moreno, P. Genomic DNA extraction and Genotyping: Ojwang, P.P.O., Njung’e, V. Compiled of the manuscript: Ojwang P.P.O and Eldridge, T.

